# Development of an on-chip fluorescence anisotropy immunoassay for human C-peptide secretion reveals a general roadmap for tracer optimization

**DOI:** 10.1101/2024.10.18.619167

**Authors:** Yufeng Wang, Nitya Gulati, Romario Regeenes, Adriana Migliorini, Amanda Oake, Maria Cristina Nostro, Jonathan V. Rocheleau

## Abstract

Fluorescence anisotropy immunoassays (FAIAs) are widely used to quantify the concentration of target proteins based on competition with a tracer in binding a monoclonal antibody. We recently designed an FAIA to measure mouse C-peptide secretion from living islets in a continuous-flow microfluidic device (InsC-chip). To develop an assay for human C-peptide, our initial selection of antibody-tracer pairings revealed the need to optimize both the dynamic range and the binding kinetics to measure the assay on-chip effectively. Here, we present strategies for developing an on-chip FAIA using two different monoclonal antibodies to achieve both a large dynamic range and high temporal resolution. The two monoclonal antibodies (Ab1 & Ab2) to human C-peptide initially showed low dynamic range and slow kinetics, preventing them from being used in an on-chip assay. To shorten the time-to-reach equilibrium for Ab1, we reengineered the tracer based on a comparison between the human and mouse C-peptide sequences, resulting in > 30-fold shorter time-to-reach equilibrium. To increase the relatively small dynamic range for Ab2, we used partial epitope mapping and targeted point mutations to increase the dynamic range by 45%. Finally, we validated both FAIAs by measuring depolarization-induced insulin secretion from individual hESC-islets in our InsC-chip. These strategies provide a general roadmap for developing FAIAs with high sensitivity and sufficiently fast kinetics to be measured in continuous-flow microfluidic devices.

## INTRODUCTION

Fluorescence anisotropy immunoassays (FAIAs) are commonly used in industrial and academic research, particularly for high-throughput screening assays ^1^. FAIAs depend on the competition between a target (e.g., protein) and a fluorophore-labelled analogue (i.e., tracer) in binding to a monoclonal antibody. The apparent fluorescence anisotropy depends on the fraction of free- and antibody-bound tracer at equilibrium and is thus inversely related to the concentration of the target analyte. Compared to other protein quantification methods, such as ELISA, FAIAs do not require multiple, long incubation steps or separation of different assay reagents, and can be easily adapted to low volumes ^2^. For this reason, FAIAs have more recently been integrated into microfluidic systems for on-chip detection of protein secretion by tissue and investigation of protein-ligand interactions ^3–6^.

The dynamic range of FAIAs is primarily determined by the size difference between the bound and unbound tracer, which can be maximized by reducing the size of the tracer peptide to the minimal epitope region ^7^. However, when developing an assay for a continuous-flow microfluidic device, the ability to use the assay on-chip will additionally be impacted by the kinetics of the reaction (i.e., time-to-reach equilibrium). Reading an FAIA before the competition reaches equilibrium artificially lowers the dynamic range. Previous studies that miniaturized FAIA onto microfluidic devices ensured the reactions reached equilibrium by increasing the length of the mixing channel (and thus the residence time) ^3,4^. However, this strategy broadens any spikes/oscillations in the signal due to dispersion and thus lowers the temporal resolution of the assay ^8^. We previously miniaturized an FAIA into a continuous-flow microfluidic device (InsC-chip) to measure the dynamic secretion of mouse C-peptide, which is co-secreted from islets in equimolar ratio with insulin ^9^. We focused on shortening the time to reach equilibrium and imaging the assay where the reaction reached >90% equilibrium. The antibody and tracer combination we selected reached equilibrium within 30 s, which allowed us to image the response immediately downstream of the loaded islets. This setup provided < 7 s temporal resolution and revealed new details in glucose-stimulated first-phase insulin release.

Here, we present strategies to optimize an assay to measure insulin secretion from individual human embryonic stem cell-derive islet-like clusters (hESC-islets) using our continuous-flow InsC-chip. However, the two antibodies available for human C-peptide showed either too slow kinetics (Ab1) or too small a dynamic range (Ab2) to be used effectively in our InsC-chip. To speed up the kinetics of Ab1 binding, we truncated the tracer to find the epitope and subsequently modified the epitope with point mutations to shorten the equilibration time ^10^. Next, we increased the dynamic range of Ab1 and Ab2 by truncating the tracer to the minimal point-mutated epitope. Finally, we used both assays to measure KCl-stimulated insulin secretion from individual hESC-islets. We envision these strategies can be more generally applied to other FAIAs for on-chip measurement with high sensitivity and temporal resolution.

## MATERIALS AND METHODS

### Human C-peptide antibodies

Antibody 1 (Ab1, GN-ID4, DSHB) was concentrated according to the vendor’s protocol by placing 4 mL antibody stock into an Amicon Ultra-4 insert (50kDa cutoff). The centrifugal filter was spun at 4000G for 10 min with flow-through discarded and the concentrate was resuspended to 4 mL with PBS. The centrifugal filter was spun again at 4000G for 10 min. The concentrate was aliquoted and stored at -20°C. Antibody 2 (Ab2, MAB80561, Biotechne) was purchased in concentrated form.

### FAIA

FAIAs were characterized in 384-well plates with glass a bottom (CLS4581, Sigma Aldrich), as previously described ^9^. Briefly, a custom-designed fluorescein-tagged C-peptide (C-peptide*) was purchased from Biomatik (Ontario, Canada). On the day of the experiment, C-peptide* was diluted in BMHH imaging buffer (125 mM NaCl, 5.7 mM KCl, 0.42 mM CaCl_2_, 0.38 mM MgCl_2_, 10 mM HEPES, and 0.1% BSA; pH 7.4) and the tracer concentration was measured by absorbance of the FITC (ε = 73000 cm^-1^M^-1^) using a DS-11+ spectrophotometer (DeNovix). For direct binding experiments, a human C-peptide monoclonal antibody (either Ab1 or Ab2) was serially diluted in C-peptide* at the indicated concentrations. For competitive binding experiments, human C-peptide (Biomatik, Canada) was serially diluted in pre-mixed labelled C-peptide* and C-peptide antibody. All assay solutions were imaged at 37°C using the custom-built wide-field fluorescence anisotropy microscope described below.

### Fabrication of continuous-flow microfluidic device

Microfluidic devices were fabricated using soft lithography as described previously, with minor modifications ^9^. Briefly, Computer-Aided Design (CAD) files of the continuous-flow microfluidic device were generated in AutoCAD (Autodesk, USA). A layer of SU-8 2075 (Micro-Chem, USA) negative photoresist with the same thickness as the device height indicated in the text was spin-coated onto 100 mm silicon wafers (University Wafer) and exposed through a master printed on transparent plastic film (CAD/Art Services, USA). PDMS-based microfluidic devices were generated from the master mold and hole-punched at inlets and outlets with a beveled 22G needle. The PDMS layers were plasma bonded to no. 1.5 glass coverslips (VWR Scientific, USA). PDMS-based solution reservoirs (∼70 μL) were plasma bonded over the device inlets and tubing was inserted into the outlets.

### Expansion and differentiation of human embryonic stem cells

The H1 human embryonic stem cell (hESC) line (WiCell, Madison, WI, USA) was cultured and expanded using Stem-MACS™iPSC-Brew XF media (Miltenyi Biotec) on tissue culture plates coated with Geltrex™ LDEV-free hESC-qualified basement membrane matrix (Gibco). The differentiation protocol began 48 hours after seeding 4-4.5×10^6^ cells per 12-well tissue culture-treated plate (Corning) and when cells were 60-90% confluent. hESC-derived islet-like cell clusters (hESC-islet) were generated with a modified version of a previously published 7-stage differentiation protocol (Misra et al., unpublished). Stage 7 hESC-islets were analyzed using flow cytometry for their efficient commitment to C-peptide^+^/NKX6.1^+^ monohormonal Δ-like cells. hESC use was approved by the Stem Cell Oversight Committee (Canadian Institute of Health Research).

### Wide-field fluorescence anisotropy microscope

Steadystate fluorescence anisotropy imaging was performed using an inverted widefield RAMM fluorescence microscope (ASI), equipped with 3 LEDs (405, 505, and 590 nm) and a stage top incubator (Okolab) ^9,11,12^. All images were collected through a 40×/0.75 NA air objective (Olympus, Richmond Hill, Canada). Incident light was linearly polarized before exciting the sample. The emission light was filtered through an eYFP (ET535/30M) emission filter, and subsequently split by an Optosplit II (Cairn, Faversham, UK) to simultaneously collect parallel (*I*_‖_) and perpendicular (*I*_⊥_) intensities on separate regions of an Iris 15 sCMOS camera (Photometrics).

### Image analysis

BMHH buffer was imaged using the same microscope settings as experimental images to determine the background. Background-corrected *I*_*‖*_ and *I*_⊥_ fluorescence intensity images were analyzed with a custom script in ImageJ to calculate pixel-by-pixel anisotropy (r) ^9,12^:

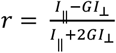

The G factor was measured by imaging a 100 nM fluorescein solution, where the equation can be simplified to G = *I*_‖_ */ I*_⊥_ ^13^.

### Fitting for FAIA

The direct and competitive binding curves of the FAIA were fitted for K_D1_ and K_D2_, as described by Roehrl *et al*. ^14^. The fluorescence anisotropy of the assay solution is related to the fraction of bound tracer (F_SB_) through ^14^:

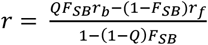

Here, r_b_ and r_f_ represent the anisotropies of bound and free tracer, respectively and Q represents the ratio of fluorescence intensity of bound to free tracer.

K_D1_ of the direct binding curve was fit in OriginLab (Northampton, MA) to the following equation:

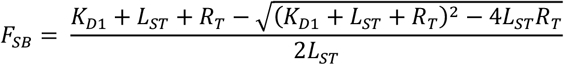

Here, L_ST_, L_T_ and R_T_ are the total concentration of the tracer, the human C-peptide, and the antibody, respectively. K_D2_ of the competitive binding curve was fit in OriginLab (Northampton, MA) to the following equations:

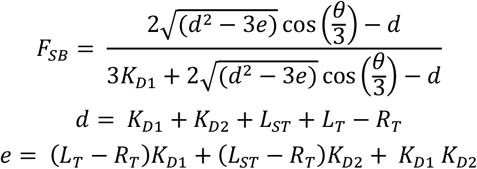

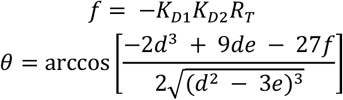

### Imaging insulin secretion

C-peptide as a surrogate for insulin secretion was measured using the InsC-chip as previously described ^9^. Briefly, hESC-islets were pre-incubated in BMHH at 1 mM glucose for 1 hr (37°C, 5% CO_2_) before loading into the InsC-chip through gravity-driven flow using height differences in the inlet and outlet tubing. The InsC-chip was mounted onto a stage-top incubator (Okolab) at 37 °C and connected to a pressure-based flow controller (Fluigent, U.S.). Treatments pre-mixed with tracer and antibody were added to the on-chip reservoir and the flow controller was set to 50 μL/hr. The pearl-shaped channel downstream of each islet trap collected effluent. This pearl shape provided a fiduciary distance marker to ensure the competition assays were imaged at the indicated distance (i.e., equilibrium time) from the islet. The relative amount of insulin secretion was reported by normalizing the anisotropy response of the tracer to that at 1 mM glucose, where insulin secretion is generally negligible.

### Statistics

Replicates are depicted as mean ± standard error of the mean (S.E.M.) if not stated otherwise. All data are based on experiments conducted on at least three separate days (n = 3 independent trials). Statistical analysis was performed using Prism 6 (GraphPad, U.S.) and the statistical tests used are specified in the text.

## Results

### Design of an on-chip FAIA to measure human C-peptide secretion

To illustrate the design criteria for on-chip FAIAs, we demonstrate the optimization of an assay for human C-peptide to measure secretion from individual hESC-islets using the InsC-chip (**Figure 1)**. The InsC-chip immobilizes 4 islets in flowing media containing an antibody-bound tracer (**Figure 1A, left**). Islets held in the separate traps secrete insulin C-peptide into separate pearl-shaped detection channels, which provide ∼100 s of on-chip residence time for the competitive binding to reach equilibrium (**Figure 1A, right**). We followed four steps to ensure the assay could be read on-chip. First, we designed fluorophore-tagged C-peptide tracers (C-peptide*) and characterized the direct binding to antibodies to fit K_D1_, the direct binding dissociation constant (Step 1, **Figure 1B**). This step was also used to raise the dynamic range of the response by truncating the tracer to the epitope region, thus minimizing the peptide size and maximizing the difference in molecular weight between the free and antibody-bound tracer (Step 1, **Figure 1B**). Second, we characterized the competitive binding between antibody-bound C-peptide* and unlabeled C-peptide to determine K_D2_, the dissociation constant for the target analyte. (Step 2, **Figure 1B**). Third, we determined the sensitivity of each FAIA based on K_D1_, K_D2_, and the concentrations of tracer and antibody ^14^. For instance, pre-mixing different concentrations of C-peptide* with a fixed antibody concentration yields distinct assay sensitivities (Step 3, **Figure 1C**). We previously confirmed that pre-mixing antibodies and tracer at concentrations that yield 50% of bound tracer (i.e., F_SB_ = 0.5) provides maximum FAIA sensitivity ^9^. It is worth noting that K_D1_ is commonly equated to the antibody concentration resulting in 50% bound protein, but it is rarely specified that this is at equilibrium, and not the total concentration of the antibody. To find the total antibody concentration ([Ab]_total_) that will yield F_SB_ = 0.5, we derived the relationship between [Ab]_total_, K_D1_ and total tracer concentration ([Tracer]_total_) (**Figure S1**). When F_SB_ = 0.5, [Ab]_total_ = K_D1_ + ½[Tracer]_total_. Finally, we evaluated the kinetics of the competition assay to determine the time-to-reach equilibrium (Step 4, **Figure 1D**). Our InsC-chip provides ∼100 s of residence time in the pearl-shaped channel to allow the assay to reach equilibrium. Modifying the chip to increase this residence time would lead to greater signal dispersion and lower the temporal resolution of the signal ^9^. Hence, our goal was to shorten the time-to-reach equilibrium to < 100 s by modifying the tracer and measuring the assay at the pearl channel where the assay reaches > 90% equilibrium.

**Figure 1.**
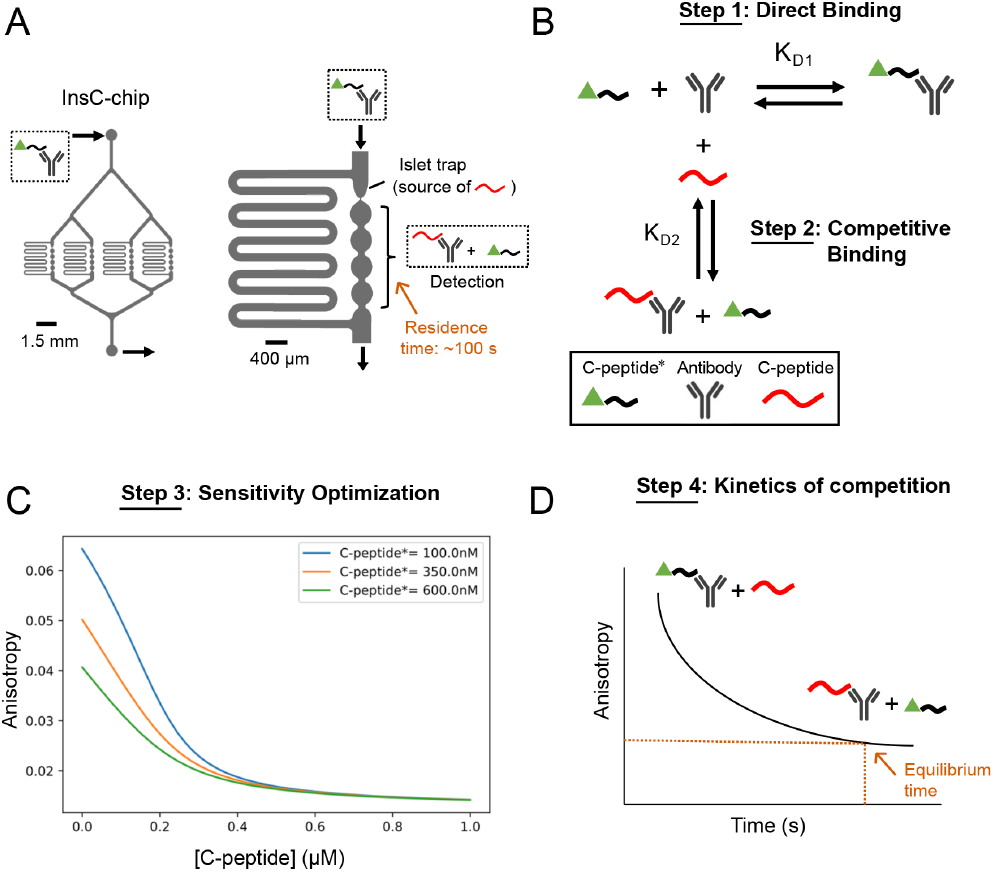
General framework for miniaturization of FAIA into microfluidics. A) Incorporation of FAIA into a microfluidic device, using the InsC-chip as an example. Pre-mixed C-peptide* and antibody is drawn through inlet of the InsC-chip, which contains four reaction chambers (left). As assay solution flows through each reaction chambers (right), C-peptide secreted from a loaded islet starts to compete with C-peptide* for antibody binding. The downstream pearl channel provides around 100 s of residence time for the FAIA to reach equilibrium on-chip. B) Schematic of the direct binding between C-peptide tracer (C-peptide*) and antibody, and the competitive binding between C-peptide* and full-length C-peptide against antibody. C) Simulation of C-peptide* with varying concentrations competing with C-peptide against a fixed concentration of antibody. D) Schematic of the temporal dynamics of the competitive binding shown in C).

### Characterization of antibody candidates for FAIA

Knowledge of the antibody epitope is critical to limit the tracer size and modulate the antibody-tracer interactions ^15,16^. Unfortunately, manufacturers do not always perform epitope mapping or consider this information proprietary. To develop an on-chip FAIA, we started with two monoclonal antibodies for human C-peptide with a stated limited affinity for mouse C-peptide. The first antibody (Ab1) had a partially mapped epitope ^17^ and the second antibody (Ab2) was previously validated for C-peptide ELISA but did not have a validated epitope. Human C-peptide comprises 31 amino acids, whereas antibody epitopes commonly contain 5-7 amino acids ^18^. To measure Ab1 and Ab2 binding, we initially designed a tracer with a fluorophore (FITC) tagged on the N-terminus of full-length human C-peptide (F-C-peptide*) (**Figure 2A**). Ab1 (**Figure 2B**) and Ab2 (**Figure 2C**) binding to F-C-peptide* showed an order of magnitude difference in K_D1_ values of 2.74 ± 1.1 and 21.1 ± 1.1 nM, respectively. To measure the affinity and kinetics of the competition assays, we serially diluted unlabeled C-peptide into premixed antibody and F-C-peptide* (**Figure 2D-G**). Ab1 showed a K_D2_ of 3.39 ± 0.21 (**Figure 2D**); however, the competition curves steadily increased in dynamic range (ΔmR) from 12.4 ± 1.05 to 43 ± 0.11 mR (milli-anisotropy), indicating the assay was still approaching equilibrium at 30 to 60 min (**Figure 2E**). In contrast, Ab2 showed a K_D2_ of 1.60 ± 0.67 nM (**Figure 2F**) and reached equilibrium within 5 min with a ΔmR of 20 ± 1.6 (**Figure 2G**). Overall, these data showed we needed to apply different strategies to optimize the tracer pairings to use these antibodies in the InsC-chip. For Ab1, we needed to modify the tracer to shorten the time-to-reach equilibrium below the ∼ 100 s residence time provided by the InsC-chip ^9^. In contrast, Ab2 showed sufficiently fast binding kinetics but a low dynamic range. Thus, our goal was to perform epitope mapping on Ab2 to design a tracer with the shortest possible peptide sequence (i.e., low molecular weight) to increase the dynamic range.

**Figure 2.**
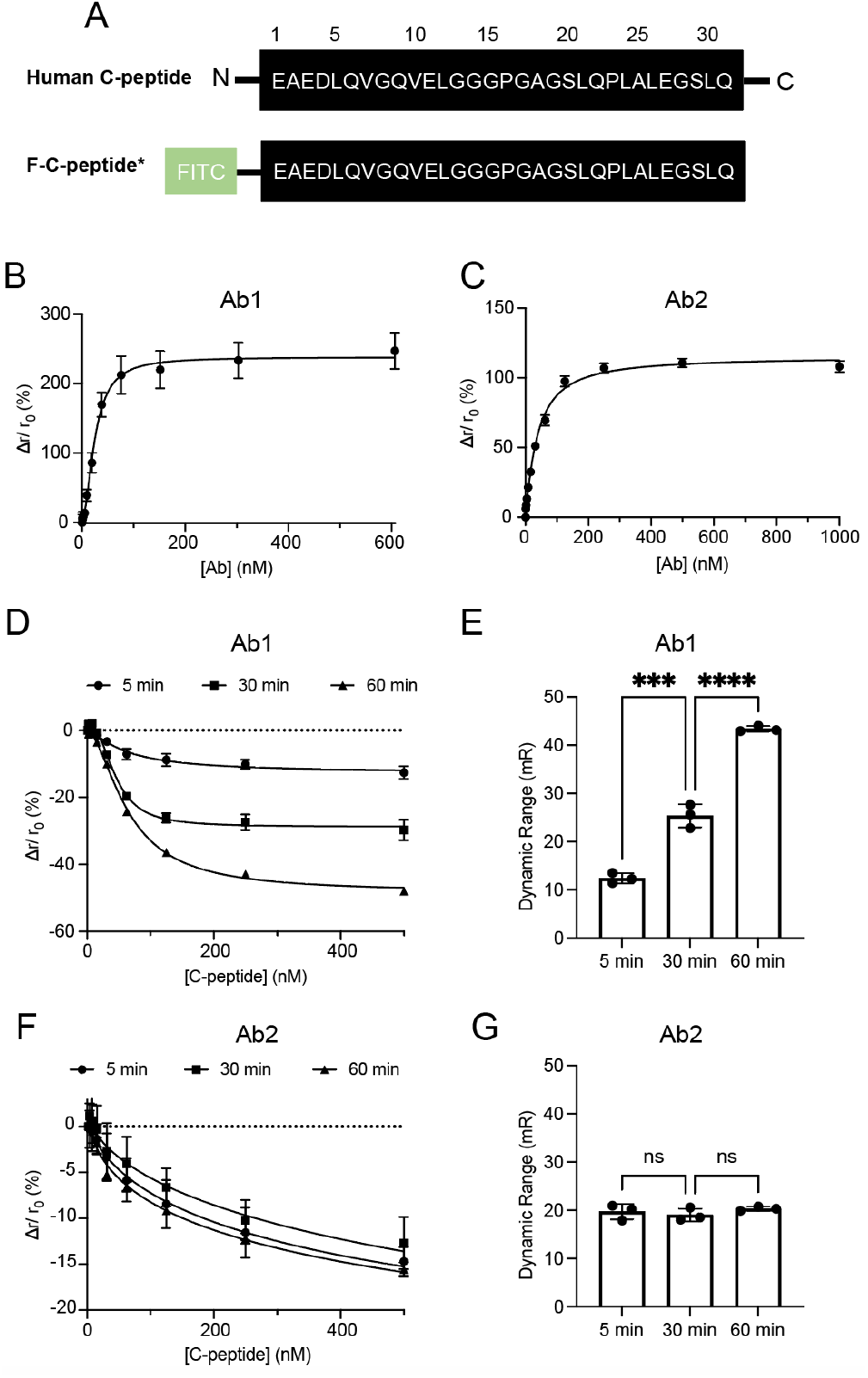
Development of F-C-peptide* for human C-peptide FAIA. A) Schematic of sequences of human endogenous C-peptide (top) and F-C-peptide* (bottom). B-C) Fitting of the direct binding curves between serially diluted Ab1 (B, n=3) or Ab2 (C, n=3) and 100 nM F-C-peptide* yielded a K_D1_ of 2.74 ± 1.1 nM or 21.1 ± 1.1 nM, respectively. D, F) Fitting of competitive binding curves between serially diluted C-peptide and pre-mixed 10 nM Ab1 + 20 nM C-peptide* or 30 nM Ab2 + 20 nM C-peptide* yielded a K_D2_ of 3.39 ± 0.21 nM (n = 3) or 1.60 ± 0.67 nM (n = 3), respectively. The assay solution was imaged after 5, 30 and 60 min of incubation in 384-well plate. E, G) Dynamic range of the competition curve for Ab1 (E) and Ab2 (G) after 5, 30 and 60 min of incubation. *** indicates p < 0.001, **** indicates p < 0.0001 by one-way ANOVA.

### Increasing FAIA kinetics through tracer point mutations

Madsen *et al*. suggested that Ab1 binds to amino acid residues 8-13 and 25-31 of human C-peptide (**Figure 3A**) ^17^. Thus, we truncated the first 7 residues of full-length C-peptide to maintain these putative epitopes in our tracers. To shorten the equilibrium time for Ab1, we looked in this shortened sequence for hydrophobic resides that primarily contribute to protein-protein interactions within epitopes ^16^, while avoiding glycine and proline, which are often critical for protein structure. We confirmed Ab1 does not cross-react with mouse C-peptide ^17^ (**Figure S2**). Thus, we also narrowed our search for point mutations by comparing the human and mouse sequences. Based on this overall strategy, we generated 3 mutant tracers (M1 – M3, **Figure 3A**).

**Figure 3.**
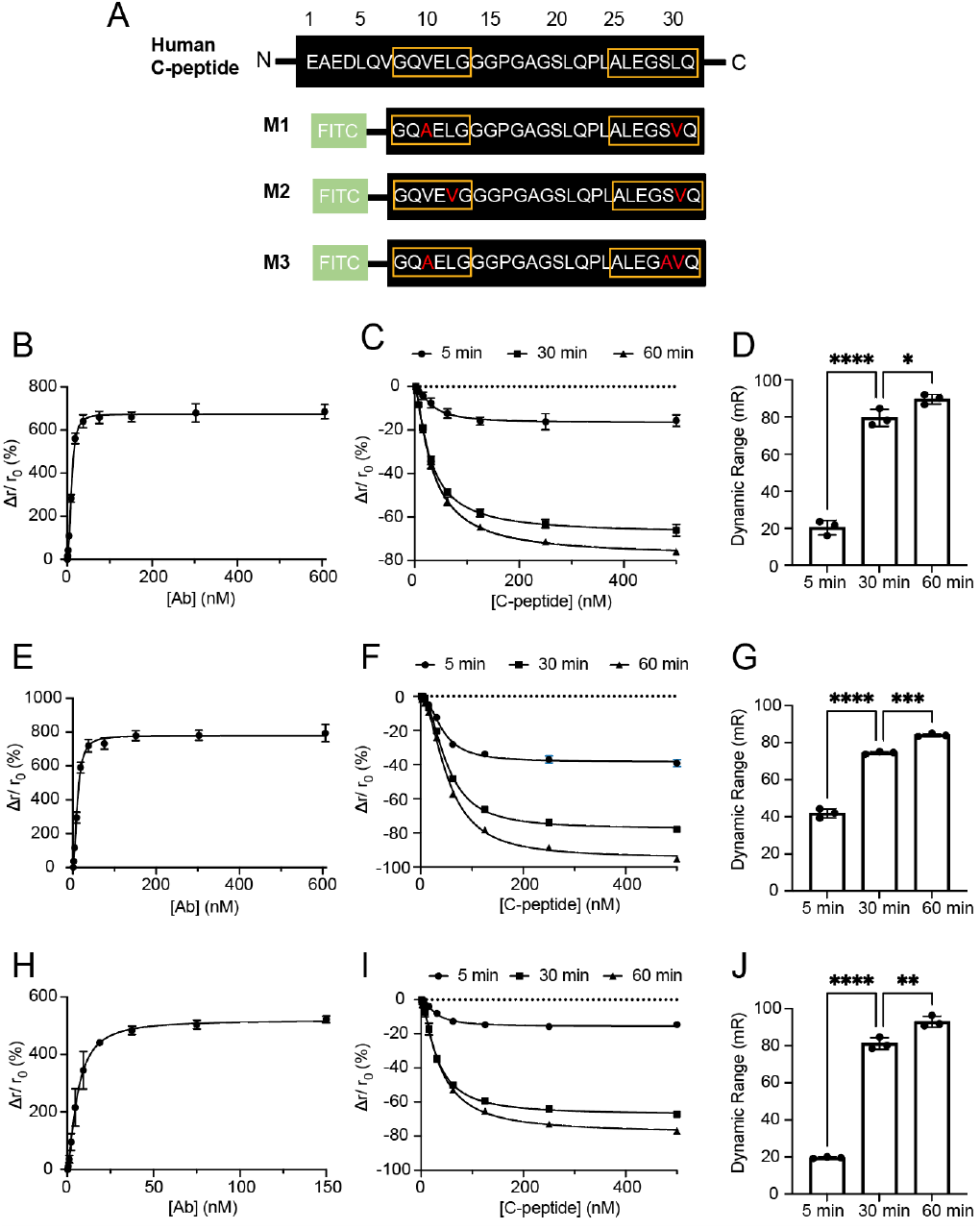
Development of M1-3 for human C-peptide FAIA with Ab1. A) Schematic of sequences of human endogenous C-peptide (top) and M1-3 (bottom). Boxed residues contain the suggested epitope region from the previous study ^17^. B, E, H) Fitting of the direct binding curves between Ab1 and 100 nM M1, 100 nM M2 or 50 nM M3 yielded a K_D1_ of 3.65 ± 0.4 nM, 4.03 ± 0.93 nM and 2.07 ± 0.43 nM, respectively (n = 3 for each tracer). C, F, I) Fitting of competitive binding curves between C-peptide and pre-mixed 10 nM Ab1 and 20 nM M1, M2 or M3 yielded a K_D2_ of 2.07 ± 1.13 nM, 3.44 ± 1.34 nM and 4.1 ± 2.2 nM (n = 3 for each tracer). The assay solution was imaged after 5, 30 and 60 min of incubation in 384-well plate. D, G, J) Dynamic range of the competition curves shown in C, F, I) after 5, 30 and 60 min of incubation. * indicates p < 0.05, ** indicates p < 0.01, *** indicates p < 0.001, **** indicates p < 0.0001 by one-way ANOVA.

In tracer mutant 1 (M1), we switched the valine and leucine at positions 10 (Val10) and 30 (Leu30) to less bulky yet still non-polar alanine and valine, respectively (**Figure 3A**). M1 showed similar Ab1 binding affinity as F-C-peptide*, with K_D1_ = 3.65 ± 0.4 nM and K_D2_ = 2.07 ± 1.13 nM (**Figure 3B-C**). The kinetics of the competition with M1 were still quite slow, as the increase in ΔmR from 20.4 ± 3.9 to 90 ± 2.7 mR took over the 1-hr incubation (**Figure 3D**). However, the competition curve for M1 after 1-hr incubation showed a significantly larger ΔmR than that for F-C-peptide*, consistent with M1 being smaller than F-C-peptide*. In tracer mutant 2 (M2), we changed Leu12, which was previously reported to be critical to antibody binding^17^ (**Figure 3A**). M2 again showed similar binding affinity to Ab1 as F-C-peptide* (K_D1_ = 4.03 ± 0.93 nM, K_D2_ = 3.44 ± 1.34 nM) with slow kinetics and a similarly increased ΔmR (84 ± 0.69 mR) at 1 hr (**Figure 3E-G**). Since mutating to smaller non-polar residues did not change the competition kinetics significantly, we subsequently mutated the polar Ser29 to Ala29 (M3), which is found in mouse C-peptide sequence (**Figure 3H-J)**. Surprisingly, such a drastic point mutation within the putative epitope region did little to alter the FAIA affinity (K_D1_ = 2.07 ± 0.43 nM, K_D2_= 4.1 ± 2.2 nM) and kinetics. Overall, mutations at positions 10, 29, and 30 only moderately changed the kinetics, suggesting the putative epitope was incorrect. Thus, for subsequent tracer design, we targeted the differences between the mouse and human sequences in the connecting region (position 14-24).

The serine at position 20 (Ser20) is the only residue within 14 to 24 segment that is different between mouse and human C-peptide, and not a proline or glycine (**Figure 4A**). To measure the role of Ser20 in binding, we generated tracer mutant (M4) by replacing the polar uncharged serine with negatively charged aspartate (S20D) found in the mouse C-peptide sequence (**Figure 4A**). Ab1 showed a significantly lower affinity for M4, as reflected in the K_D1_ value (1060 ± 93 nM) (**Figure 4B**). We could not perform a competitive binding assay with M4 due to the K_D1_ value being close to the stock concentration of Ab1. Nonetheless, these data show Ser20 is within the epitope and critical in the C-peptide-Ab1 interaction. To moderate the impact on antibody binding, we generated tracer mutant 5 (M5) by replacing Ser20 with similarly charged yet slightly larger threonine (S20T, **Figure 4A**). Compared to F-C-peptide*, M5 showed ∼ 10 times higher K_D1_ (25.6 ± 3.9 nM) and K_D2_ (28 ± 2.6 nM) with Ab1 (**Figure 4C-D**). The equilibrium was also reached within 5 min (**Figure 4E**). However, the competitive binding showed a comparatively small ΔmR (32 ± 0.13 mR) (**Figure 4E**), suggesting the increase in FAIA kinetics came at cost of the dynamic range. To more accurately measure the equilibrium time, we used a continuous-flow microfluidic device (**Figure S3, Figure 4F-G**) ^9^. These data show competitive binding to Ab1 with M5 tracer reaches > 90% equilibrium by ∼ 65 s at varying C-peptide concentrations (**Figure 4F-G**).

**Figure 4.**
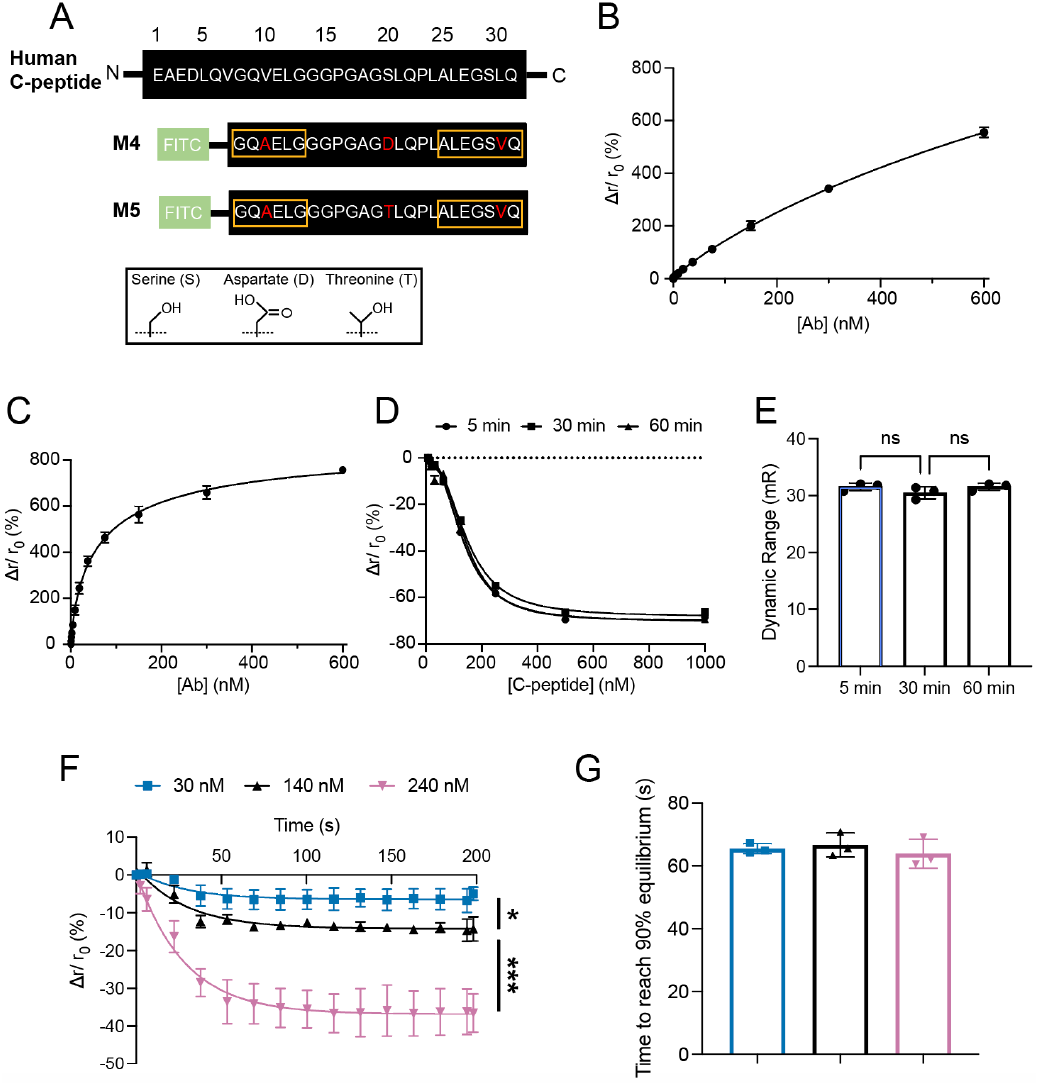
Development of M4-5 for human C-peptide FAIA with Ab1. A) Schematic of sequences of human endogenous C-peptide (top) and M4-5 (bottom). Boxed residues contain the suggested epitope region from the previous study ^17^. The insert shows the R group of serine, aspartate and threonine. B-C) Fitting of the direct binding curves between Ab1 and 50 nM M4 or M5 yielded a K_D1_ of 1060 ± 93 nM and 25.6 ± 3.9 nM, respectively (n = 3 for each tracer). D) Fitting of competitive binding curves between serially diluted C-peptide and premixed 55 nM Ab1 and 60 nM M5 yielded a K_D2_ of 28 ± 2.6 nM (n = 3) and dynamic range of 32 ± 0.13 mR. The assay solution was imaged after 5, 30 and 60 min of incubation in 384-well plate. E) Dynamic range of the competition curves shown in D) after 5, 30 and 60 min of incubation. Analysis was done using one-way ANOVA. F) Kinetics of competitive binding between different concentrations of C-peptide and pre-mixed 55 nM Ab1 and 60 nM M5 in the device shown in **Figure S3** (n = 3 for each C-peptide concentration). Anisotropy values after 70 s of competition with the indicated concentration of C-peptide were used for statistical analysis. * indicates p < 0.05, *** indicates p = 0.0006 by one-way ANOVA. G) Time taken to reach 90% plateau of the competition kinetics curves shown in F).

### Increasing FAIA dynamic range through tracer truncation

To increase the dynamic range of Ab1 and M5, we truncated both termini as they were likely not critical in binding to Ab1 (**Figure 5**). Madsen *et al*. showed that truncating to Leu12 significantly impacted peptide-Ab1 interaction ^17^. Hence, mutant tracer M6 containing the critical S20T mutation was based on residues 12 to 24 (**Figure 5A**). M6 showed a lower affinity than M5 for binding to Ab1 with a K_D1_ of 42 ± 3.6 nM and K_D2_ 32 ± 0.36 nM (**Figure 5B-D**); yet the ΔmR of the competition curve increased from 32 ± 0.13 (M5) to 82 ± 0.93 mR (M6) (**Figure 5C-D**). Critically, the kinetics of the competition involving M5 and M6 were similar (**Figure 5E-F**), requiring < 80 s to reach > 90% equilibrium. At this equilibrium time, the FAIA using Ab1 and M6 could be measured at approximately the 4^th^ pearl of the InsC-chip.

**Figure 5.**
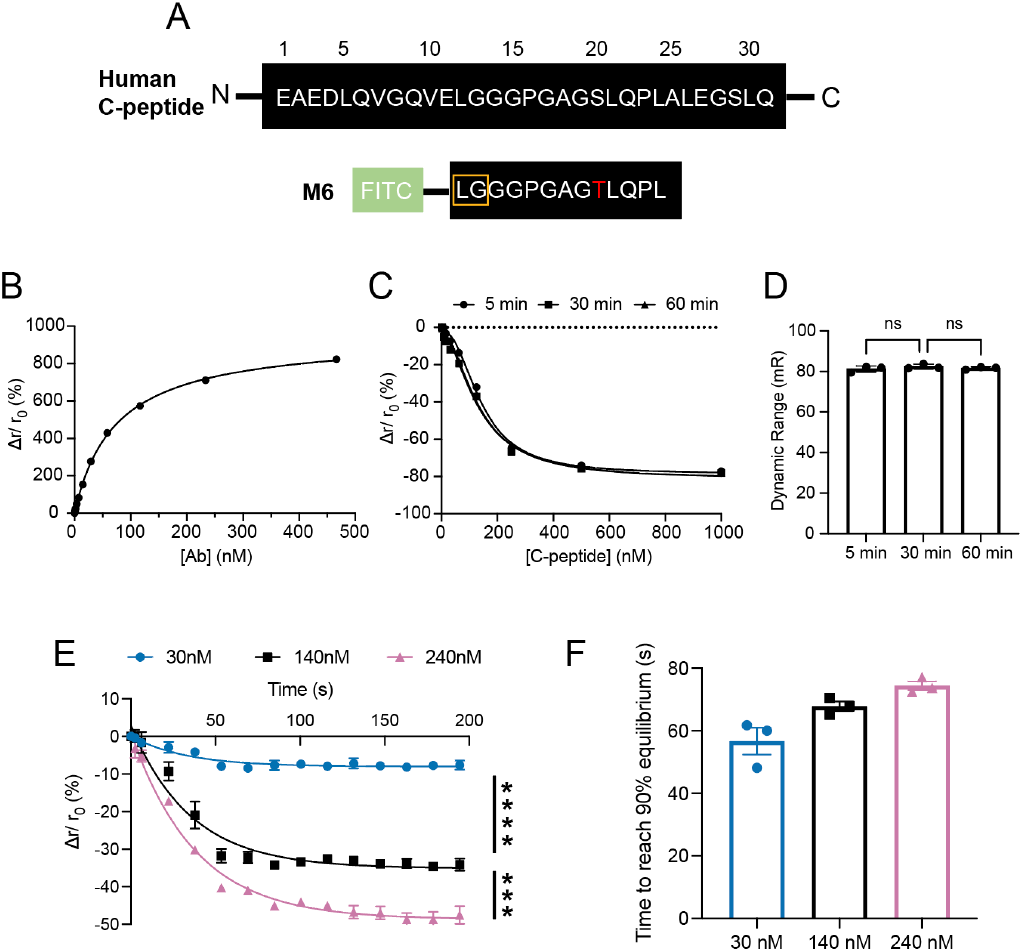
Development of M6 for human C-peptide FAIA with Ab1. A) Schematic of sequences of human endogenous C-peptide (top) and M6 (bottom). Boxed residues contain the suggested epitope region from the previous study ^17^. B) Fitting of the direct binding curves between Ab1 and 50 nM M6 yielded a K_D1_ of 42 ± 3.6 nM (n = 3). C) Fitting of competitive binding curves between serially diluted C-peptide and pre-mixed 70 nM Ab1 and 60 nM M6 yielded a K_D2_ of 32 ± 0.36 nM (n = 3) and dynamic range of 82 ± 0.93 mR. The assay solution was imaged after 5, 30 and 60 min of incubation in 384-well plate. D) Dynamic range of the competition curves shown in C) after 5, 30 and 60 min of incubation. Analysis was done using one-way ANOVA. E) Kinetics of competitive binding between different concentrations of C-peptide and pre-mixed 70 nM Ab1 and 60 nM M6 in the device shown in **Figure S3** (n = 3 for each C-peptide concentration). Anisotropy values after 80 s of competition with the indicated concentration of C-peptide were used for statistical analysis. *** indicates p < 0.001, **** indicates p < 0.0001 by one-way ANOVA. F) Time taken to reach 90% plateau of the competition kinetics curves shown in E).

As mentioned previously, Ab2 showed sufficiently fast kinetics but a low dynamic range. A prominent strategy to increase the dynamic range of an FAIA is to lower the tracer size to below 1500 Da or ∼13 amino acids ^19^. Thus, our strategy to find the epitope region of Ab2 was to divide human C-peptide (31 residues) into pieces of 10 to 13 residues. We first designed mutant tracer 7 (M7) by truncating full-length C-peptide from both ends, keeping residues from position 12 to 24 (**Figure 6A**). Compared to F-C-peptide*, M7 showed elevated K_D1_ (58 ± 1.1 nM) and K_D2_ (110 ± 10 nM) (**Figure 6B-C**). Critically, ΔmR of the competitive binding almost tripled to 58 ± 2.8 mR compared to F-C-peptide* (**Figure 6D**). These data suggest that the epitope for Ab2 is contained within residues 12 to 24. Next, we measured the binding kinetics in the continuous-flow microfluidic device (**Figure S4A-B**). These data showed it took < 20 s for the competition to reach 90% equilibrium (**Figure S4A-B**).

**Figure 6.**
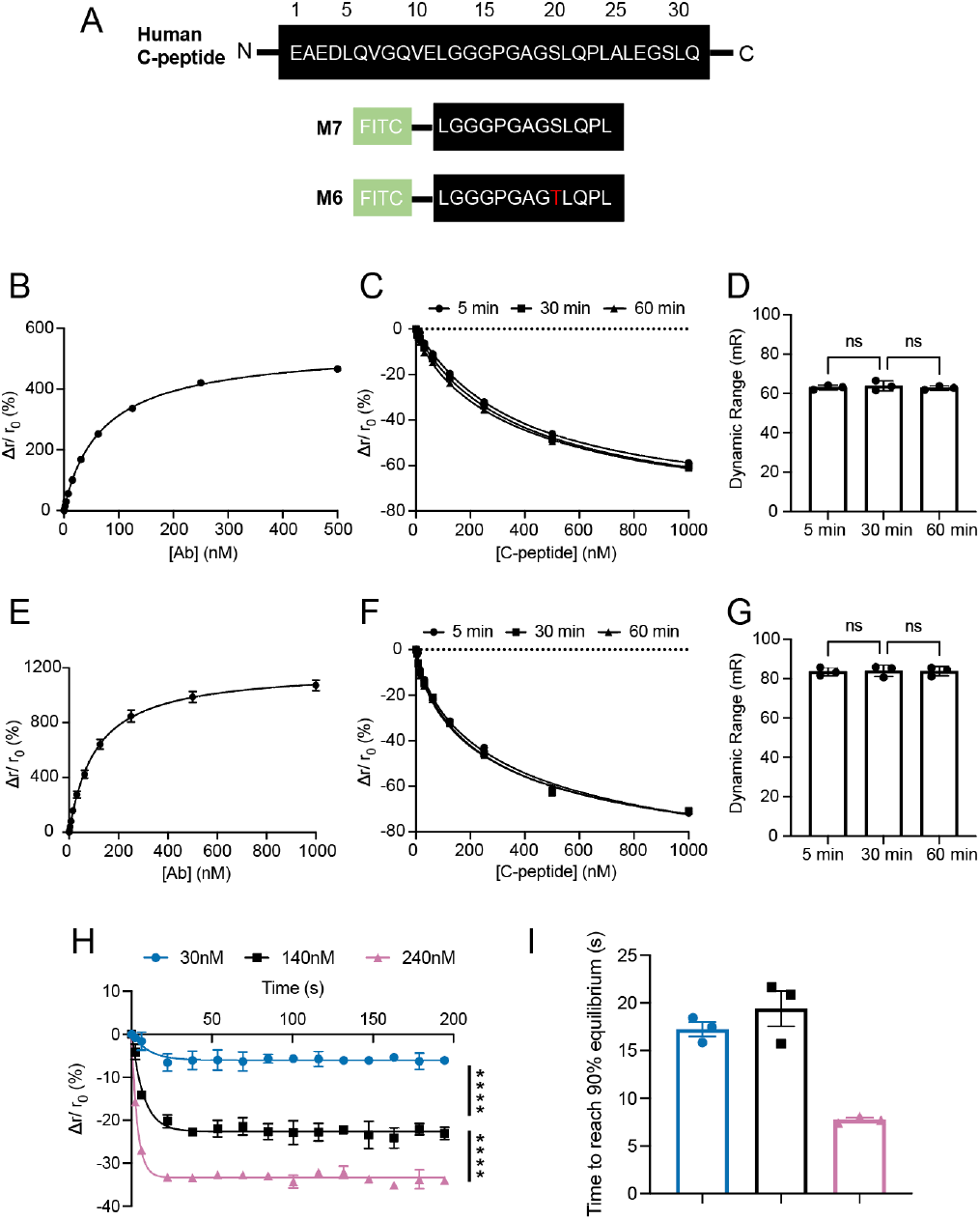
Development of C-peptide* for human C-peptide FAIA with Ab2. A) Schematic of sequences of human endogenous C-peptide (top) and M6-7 (bottom). B, E) Fitting of the direct binding curves between Ab2 and 50 nM M7 or M6 yielded a K_D1_ of 58 ± 1.1 and 70 ± 3.7 nM, respectively (n = 3 for each tracer). C, F) Fitting of competitive binding curves between serially diluted C-peptide and pre-mixed 90 nM Ab2 + 60 nM M7 or 100 nM Ab2 + 60 nM M6 yielded a K_D2_ of 110 ± 10 or 77 ± 3.7 nM (n = 3 for each tracer) and dynamic range of 58 ± 2.8 or 84 ± 2.1 mR. The assay solution was imaged after 5, 30 and 60 min of incubation in 384-well plate. D, G) Dynamic range of the competition curves shown in C, F) after 5, 30 and 60 min of incubation. Analysis was done using one-way ANOVA. H) Kinetics of competitive binding between different concentrations of C-peptide and pre-mixed 100 nM Ab2 and 60 nM M6 in the device shown in **Figure S3** (n = 3 for each C-peptide concentration). Anisotropy values after 22 s of competition with the indicated concentration of C-peptide were used for statistical analysis. **** indicates p < 0.0001 by one-way ANOVA. I) Time taken to reach 90% plateau of the competition kinetics curves shown in H).

To improve the dynamic range of the FAIA using Ab2, we applied a similar tracer design previously used for Ab1 (**Figure 6**). Based on the manufacturer, Ab2 is specific to human C-peptide. To guide our selection of mutations, we confirmed Ab2 does not cross-react with mouse C-peptide (**Figure S5**). Again, we avoided glycine and proline residues that differ between human and mouse C-peptide. This comparison revealed tracer M6 was an ideal candidate for Ab2, similar to Ab1 (**Figure 6A**). M6 showed similar binding characteristics as M7 in binding Ab2, with K_D1_ of 70 ± 3.7 nM and K_D2_ of 77 ± 3.7 nM (**Figure 6E-F**). Interestingly, the S20T point mutation in M6 significantly increased ΔmR of the competition curve from 58 ± 2.8 (M7) to 84 ± 2.1 mR (M6) (**Figure 6G**). The kinetics of the competition were also similar between the M7 and M6 (**Figure 6H-I**), requiring only ∼ 20 s to reach > 90% equilibrium. Thus, an FAIA using Ab2 and M6 tracer can be measured at the 1^st^ pearl in our InsC-chip. Overall, these data suggest that both peptide truncation and point mutation could modulate the dynamic range of FAIA.

### On-chip FAIA validation in hESC-islet

M6 binding by Ab1 and Ab2 showed ∼ 80 and ∼ 20 s to reach >90% equilibrium, respectively (**Figure 5F, 6I**), both sufficiently fast for miniaturization into our InsC-chip. The two antibodies show similar dynamic ranges at equilibrium (**Figure S6A**) while Ab1 showed a slightly higher sensitivity than Ab2 (**Figure S6B**). To directly compare Ab1 and Ab2 for on-chip FAIA, we measured KCl-stimulated insulin secretion from hESC-islets using each antibody and M6 tracer (**Figure 7**). Before *in vivo* implantation, hESC-islets show limited glucose-stimulated insulin secretion ^20,21^, yet secrete insulin in response to membrane depolarization by KCl stimulation. Thus, hESC-islets were treated with 30 mM KCl to image secretion using each antibody but separated by a 30-min incubation at 1 mM glucose to induce a resting state (**Figure 7A**). Each length of pearl in the mixing channels provides about ∼25 s residence time ^9^. To determine the impact of matching equilibration time to residence time on the responses, we imaged the fluorescence anisotropy at the 1^st^ and 4^th^ pearl for both antibodies (**Figure 7B-F**). A representative trace collected from the same hESC-islet at the 1^st^ pearl shows a burst of insulin release in response to 30 mM KCl, with a higher overall response using Ab2 (**Figure 7C**). The normalized area under the curve (AUC) for many hESC-islets confirmed that the responses measured using Ab2 were significantly larger than when using Ab1 (**Figure 7D**). These data are consistent with Ab2 reaching equilibrium by the 1^st^ pearl, whereas Ab1 did not (**Figure 7D**). The same spike of insulin secretion from the same hESC-islet was imaged at the 4^th^ pearl (**Figure 7E**). At this position, the curves were inverted with Ab1 showing a higher peak height than Ab2, consistent with the higher sensitivity of Ab1 compared to Ab2 (**Figure 7E**). The AUC for the Ab1 traces also increased from the 1^st^ to the 4^th^ pearl (**Figure 7F**). These data confirm the Ab1 assay had not reached equilibrium by the 1^st^ pearl. In contrast, the maximum height (**Figure 7E**) and AUC (**Figure 7F**) for Ab2 were slightly diminished at the 4^th^ pearl, consistent with signal dispersion after reaching equilibrium at the 1^st^ pearl. Overall, these data show both antibodies can be used with M6 tracer to measure insulin secretion of hESC-islets using the InsC-chip.

**Figure 7.**
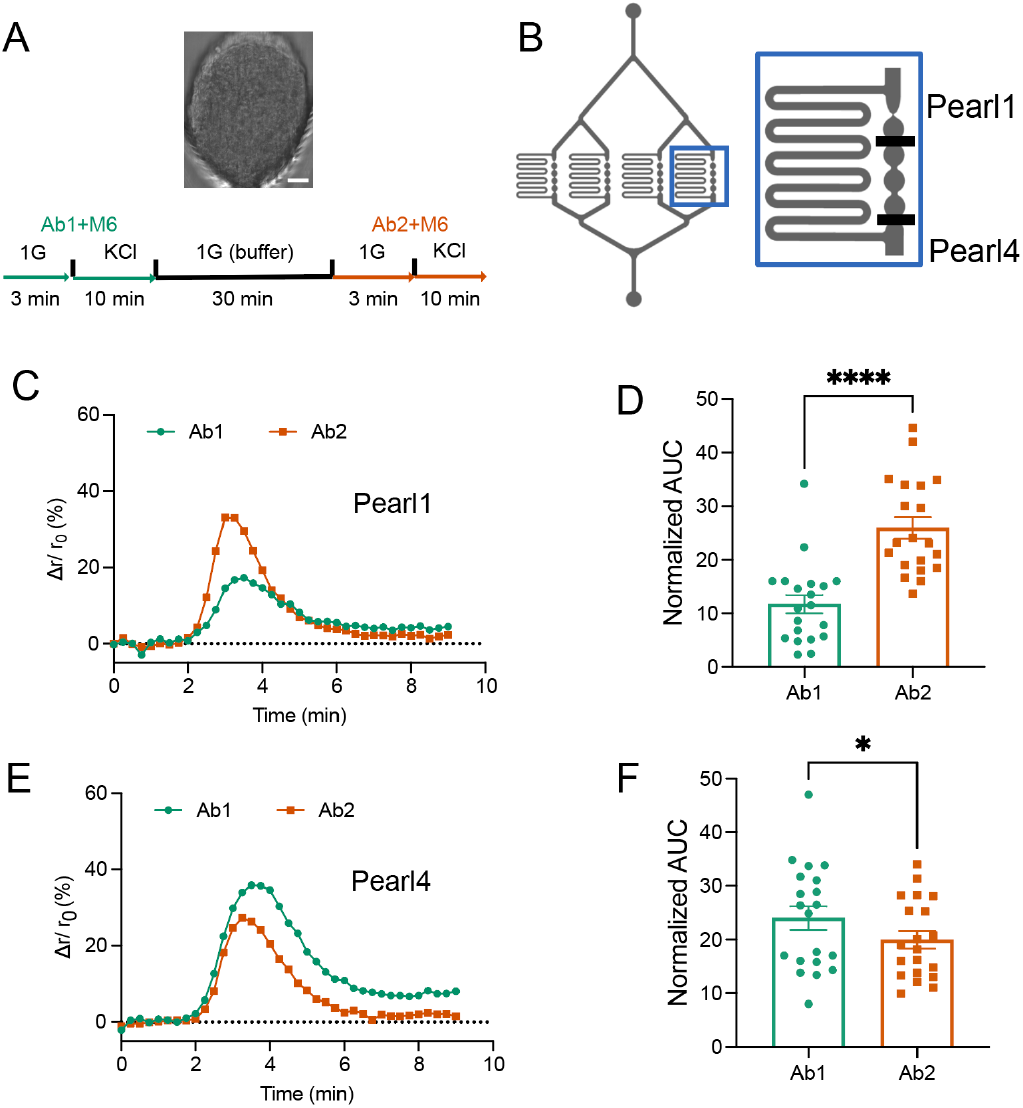
Comparison of Ab1 and Ab2 in measuring KCl-stimulated insulin secretion from hESC-islets. A) Schematic of the experiment setup. hESC-islets loaded into the InsC-chip (top, brightfield image) were treated with 1 mM glucose followed by 30 mM KCl. Secretion was measured with Ab1 and M6. 1 mM glucose in BMHH buffer were flowed through the device with the loaded SC-islets for 30 min. The same treatments were used and secretion was measured using Ab2 and M6. Scale bar represents 25 μm. B) Schematic of the InsC-chip (left) and one of the four identical reaction chambers (right). For each loaded hESC-islet, secretion was measured from Pearl 1 and 4 simultaneously. C, E) Representative traces of KCl-stimulated insulin secretion from the same SC-islet measured with Ab1 or Ab2 + M6 at Pearl 1 (C) or Pearl 4 (E). Individual traces were aligned at the initiation of secretion for comparison (n = 15). The change in anisotropy of C-peptide* (Δr) was normalized to the anisotropy just prior to KCl arrival (r_0_). D, F) Pooled area under the curve (AUC) of KCl-stimulated insulin secretion shown in C) and E) measured with different Ab-tracer combination at Pearl 1 and 4. * indicated p < 0.05, **** indicates p < 0.0001 by paired t-test.

## DISCUSSION

Fluorescence anisotropy is a sensitive and robust method to study protein-protein interactions ^2,22–24^. Methods based on fluorescence anisotropy, such as FAIA, have recently been incorporated into microfluidics to study protein secretion from living tissues or protein-protein interactions on-chip ^3–5^. We previously designed an FAIA to measure C-peptide secretion from mouse islets using an islet-on-a-chip device (InsC-chip) ^9^. Here, we modified the FAIA to measure human C-peptide secretion from hESC-islets using two monoclonal antibodies (Ab1 and Ab2) that showed intrinsically different binding affinity to C-peptide. Through the design of 6 mutant tracers for Ab1 (**Table S1**) and 2 mutant tracers for Ab2 (**Table S2**), we reveal a general roadmap to design the tracer based on enhancing dynamic range and based on speeding up the binding kinetics to measure the response on-chip. We subsequently validated both antibodies in our InsC-chip by measuring human insulin secretion from individual hESC-islets.

FAIA kinetics is a critical design factor for continuous-flow devices as it determines both the sensitivity and temporal resolution of the on-chip readout. Previous studies resorted to long mixing/ reaction channels to ensure assay equilibrium ^4,25^. However, this could lead to signal dispersion. We instead focused on modifying the FAIA kinetics. Antibody epitopes contain hot spots that contribute significantly to the protein-protein binding energy ^16^. Here, we revealed point mutations within the epitope can effectively modulate the kinetics of the FAIA. Although the epitope is rarely determined/ disclosed by the manufacturers, species specificity is often reported. We confirmed that Ab1 and Ab2 were both specific to human and not mouse C-peptide, suggesting the residues that differ between the two species contain the hot spots and are ideal targets for point mutation to control FAIA kinetics. This strategy unveiled Ser20 as a hot spot for Ab1-C-peptide since the S20D mutation (M4) decreased Ab1 binding affinity by 400x. In contrast, mutations at positions 10, 29, and 30 (M1-3), which also differ between mouse and human C-peptide, had little impact on the binding affinity. These residues may still contribute to the stabilization of antibody-protein binding, as mutations at these positions slightly increased the kinetics of competitive binding. Finally, the S20T mutation in the M6 tracer showed a nearly 10x increase in K_D1_ accompanied by kinetics acceleration by > 30×. Overall, point mutations to the tracer based on differences between the mouse and human sequences revealed a critical epitope hot spot in the antibody-protein binding. These critical amino acids serve as an ideal site for controlling FAIA kinetics. In the case of Ab1, we were able to shorten the time-to-reach equilibrium for the assay to be imaged on-chip (< 100 s).

At equilibrium, FAIA with a larger dynamic range can measure analyte with higher sensitivity. The dynamic range is primarily determined by the difference in molecular size between the bound and unbound tracer. One way to increase this difference is to lower the size of the unbound tracer ^7^. Our data confirmed this strategy as truncating non-essential residues from the tracer significantly improved the dynamic range. Examples of this strategy include the truncation of 7 residues from full-length C-peptide to generate M1 for Ab1 and the truncation of 18 residues to generate M7 for Ab2. Importantly, we demonstrated that point mutation within the epitope can also modulate the dynamic range. The S20T point mutation (M5) reduced the dynamic range of the FAIA threefold for Ab1 binding. However, the same point mutation (S20T) increased the dynamic range for Ab2 binding to M6 by 45% compared to M7 (**Table S2**). These changes in the dynamic range likely reflect the effect of short-range protein-protein interactions, such as hydrophobic and dipole-dipole forces, on k_off_ (i.e., rate of dissociation) between tracer and antibody ^10^. The value of k_off_ influences the proportion of free tracers at any given moment ^26^. A higher k_off_ results in a greater percentage of free tracer when measuring steady-state fluorescence anisotropy even at equilibrium, leading to a smaller change in fluorescence anisotropy between the bound and unbound states of the tracer. Hence, it is likely that the same S20T mutation impacted the k_off_ for Ab1 and M5 differently than for Ab2 and M6. For example, the decrease in dynamic range between Ab1 with M5 tracer might be caused by an increase in k_off_. In contrast, the enhancement in dynamic range between Ab2 and M6 might be caused by a decrease in k_off_.

When miniaturizing an FAIA with optimized kinetics and dynamic range into a microfluidic device, the assay’s equilibrium time should match the on-chip residence time to minimize signal dispersion. To illustrate the importance of matching the two time-scales, we measured KCl-induced secretion from hESC-islets using the InsC-chip, which provides a ∼ 100 s maximum equilibrium time. The time-to-reach equilibrium for Ab1 with the M6 tracer was < 80 s. We measured responses using this pairing at the 1^st^ pearl, where the tracer response was small as the assay had not yet reached equilibrium. Measuring the same secretion using the same pairing at the 4^th^ pearl provided adequate time for the assay to reach equilibrium, which yielded a larger tracer response. Ab2 with M6 tracer, in contrast, requires < 20 s to reach equilibrium. This pairing showed diminished responses from the 1^st^ to 4^th^ pearl of the InsC-chip. This is consistent with increased signal dispersion when providing a residence time longer than required to reach equilibrium. Hence, our data highlights the added constraint of optimizing the FAIA kinetics for miniaturization into a microfluidic device.

## CONCLUSION AND LIMITATION

We demonstrated strategies for optimizing the dynamic range and kinetics of FAIA assays using two antibody candidates to measure human insulin secretion from individual hESC-islets using the InsC-chip. Both antibodies (Ab1 and Ab2) showed very different binding affinities and kinetics to full-length human C-peptide. To speed up the kinetics of the FAIA using Ab1, we introduced point mutations into the tracer peptide based on differences between mouse and human C-peptide sequences. This shortened the equilibrium time for Ab1 by > 30×. This strategy also helped maximize the dynamic range of FAIA using Ab2 by truncating the tracer peptide down to a point-mutated epitope region, resulting in a fourfold enhancement of the assay dynamic range.

Our strategy led to two FAIAs for human C-peptide with different characteristics. However, our optimization strategies do have limitations. First, the generation of mutant tracers in the absence of a known epitope can be an expensive process. Second, our method relied heavily on the differences in protein sequence between species. For proteins with long and/ or divergent sequences, a larger library of point mutants will be required to find the residue(s) critical for assay kinetics. For example, human IFN-Δ is 166 amino acids long and only shares ∼50% sequence similarity with mouse ^27^. Finally, some protein sequences, such as glucagon, are identical among many mammals. In this case, one may need many more mutations to properly map the epitope, thus significantly increasing the cost and time to find point mutations that provide optimal kinetics and dynamic range.

## ASSOCIATED CONTENT

### Supporting Information

The Supporting Information is available free of charge on the ACS Publications website.

Relationship among K_D1_, total concentration of antibody and tracer (Figure S1); Ab1 does not cross-react with mouse C-peptide (Figure S2); continuous-flow microfluidic device for measuring FAIA kinetics (Figure S3); kinetics of the competitive binding for Ab2 and M7 (Figure S4); Ab2 does not cross-react with mouse C-peptide (Figure S5); comparison of competitive binding for Ab1 and Ab2 with M6 (Figure S6); summary of C-peptide* developed for Ab1 (Table S1); summary of C-peptide* developed for Ab2 (Table S2); (PDF)

## AUTHOR INFORMATION

### Author Contributions

Y.W. and J.V.R. designed the research; Y.W., N.G., and R.R. performed the research. Y.W. analyzed data; A.O., A.M. and C.N. provided hESC-islets. Y.W. and J.V.R. wrote the manuscript.

### Notes

The authors declare no competing interests.

## ACKNOWLEDGMENT

This work was supported by grants from NSERC (RGPIN-2022-04454 and DGDND-2022-04454) to J.V.R. Stipend support was provided by to Y.W by JDRF Canada through the Canadian Islet Research and Training Network (CIRTN). A.O. was supported through Breakthrough T1D International (made possible through collaboration between Breakthrough T1D International and the Leona M. and Harry B. Helmsley Charitable Trust), the Banting and Best Diabetes Centre (funded by Eli Lilly Canada), and NSERC-CREATE through CIRTN.

## Notes

### Competing Interest Statement

The authors have declared no competing interest.

