## Supplemental File for "Development of an on-chip fluorescence anisotropy immunoassay for human C-peptide secretion reveals a general roadmap for tracer optimization"

$$\text{Ab-Tracer} \xleftrightarrow{K_{D1}} \text{Ab} + \text{Tracer}$$

By definition :

$$\begin{cases} K_{D1} = \dfrac{[\text{Ab}][\text{Tracer}]}{[\text{Ab}-\text{Tracer}]} \\ F_{SB} = \dfrac{[\text{Tracer}]_{\text{total}} - [\text{Tracer}]}{[\text{Tracer}]_{\text{total}}} = \dfrac{[\text{Ab}-\text{Tracer}]}{[\text{Tracer}]_{\text{total}}} \\ [\text{Ab}]_{\text{total}} = [\text{Ab}] + [\text{Ab\_Tracer}] \end{cases}$$

When $F_{SB} = 0.5$ :

$$\begin{cases} \frac{1}{2}[\text{Tracer}]_{\text{total}} = [\text{Tracer}] = [\text{Ab-Tracer}] \\ K_{D1} = [\text{Ab}] \\ \boxed{[\text{Ab}]_{\text{total}} = K_{D1} + \frac{1}{2}[\text{Tracer}]_{\text{total}}} \end{cases}$$

**Figure S1. Relationship among $K_{D1}$, total concentration of antibody and tracer.** Derivation of the relationship among total Ab concentration ($[\text{Ab}]_{\text{total}}$), $K_{D1}$ and total tracer concentration ($[\text{Tracer}]_{\text{total}}$). [Ab], [Tracer] and [Ab-Tracer] represents the concentration of free antibody, tracer and antibody-tracer complex. The derivation assumes 1:1 binding between antibody and tracer.

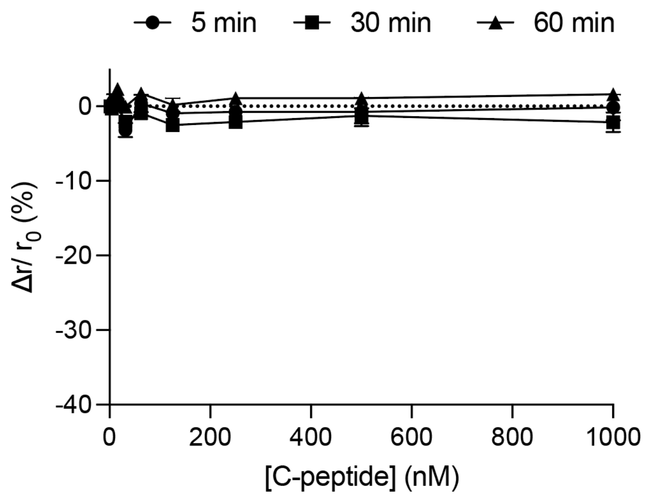


**Figure S2. Ab1 does not cross-react with mouse C-peptide.** Competitive binding curves between serially diluted mouse C-peptide and pre-mixed 10 nM Ab1 + 20 nM F-C-peptide*. The assay solution was imaged after 5, 30 and 60 min of incubation in 384-well plate.

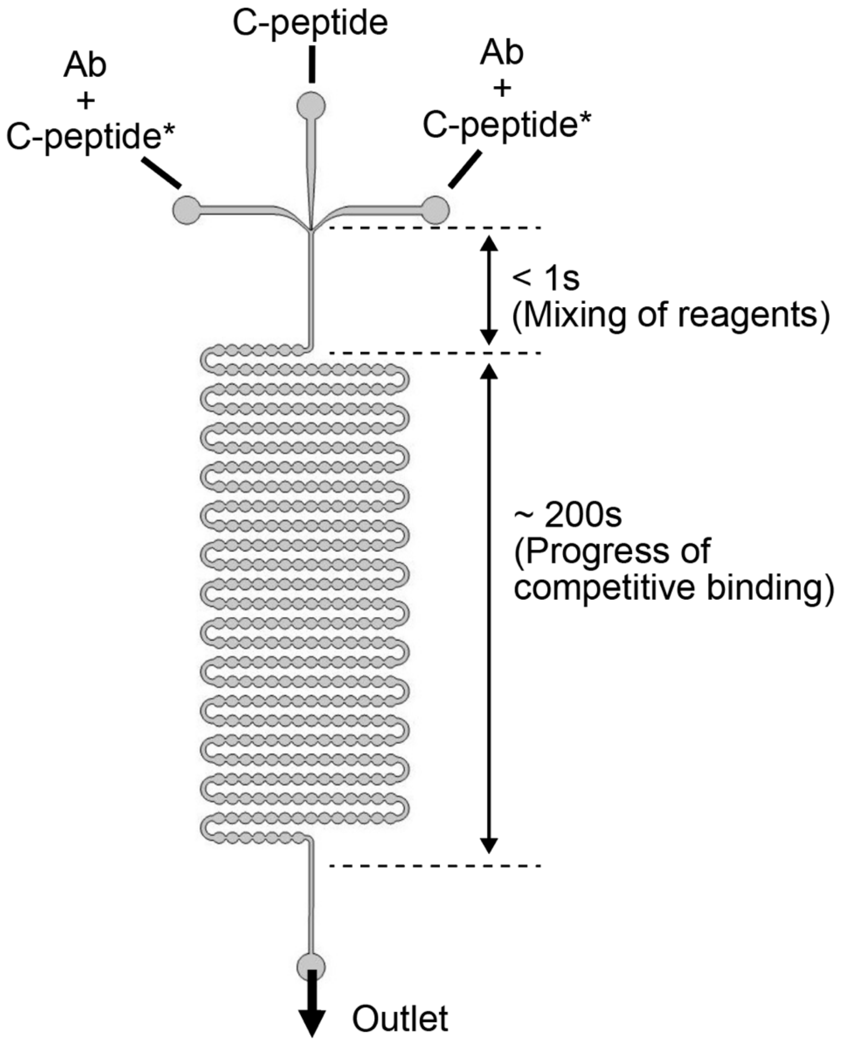


**Figure S3. Continuous-flow microfluidic device for measuring FAIA kinetics.** Schematic of the continuous-flow device for measuring the temporal dynamics of FAIA. Full-length C-peptide was flowed through the central inlet, sandwiched by premixed antibody (Ab)-C-peptide* solution in the outer two inlets. The initial straight channel allows mixing of assay reagents within 1 s. The series of pearl channels downstream provides around 200 s of residence time for the competitive binding to complete on-chip.

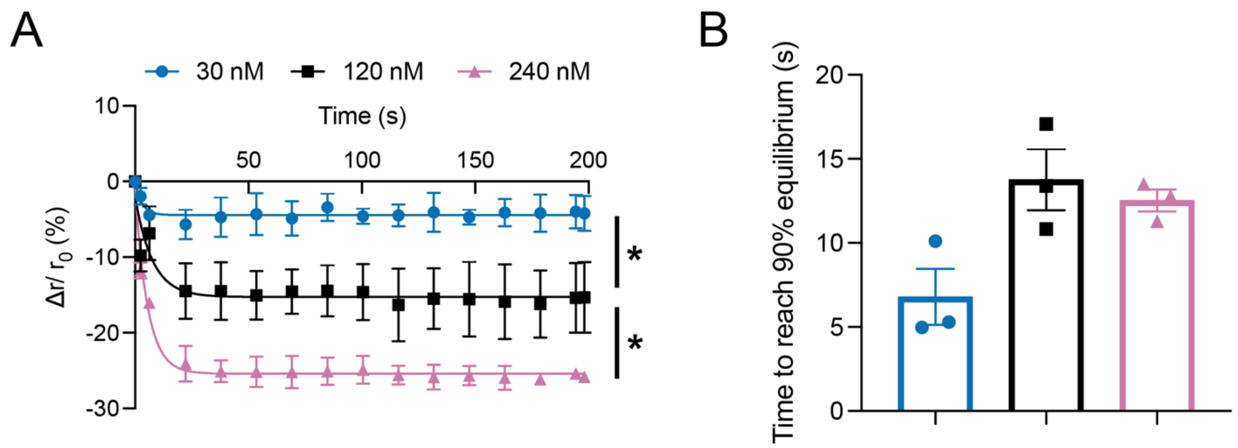


**Figure S4. Kinetics of the competitive binding for Ab2 and M7.** A) Kinetics of competitive binding between different concentrations of C-peptide and pre-mixed 90 nM Ab2 + 60 nM M7 in the device shown in **Figure S3** (n = 3 for each C-peptide concentration). Anisotropy values after 22 s of competition with the indicated concentration of C-peptide were used for statistical analysis. * indicates $p < 0.05$ by one-way ANOVA. B) Time taken to reach 90% plateau of the competition kinetics curves shown in A).

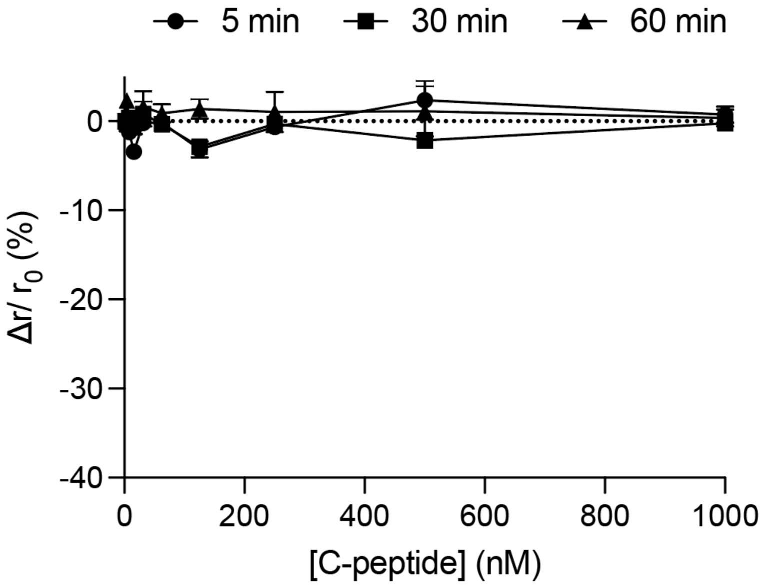


**Figure S5. Ab2 does not cross-react with mouse C-peptide.** Competitive binding curves between serially diluted mouse C-peptide and pre-mixed 30 nM Ab2 + 20 nM F-C-peptide*. The assay solution was imaged after 5, 30 and 60 min of incubation in 384-well plate.

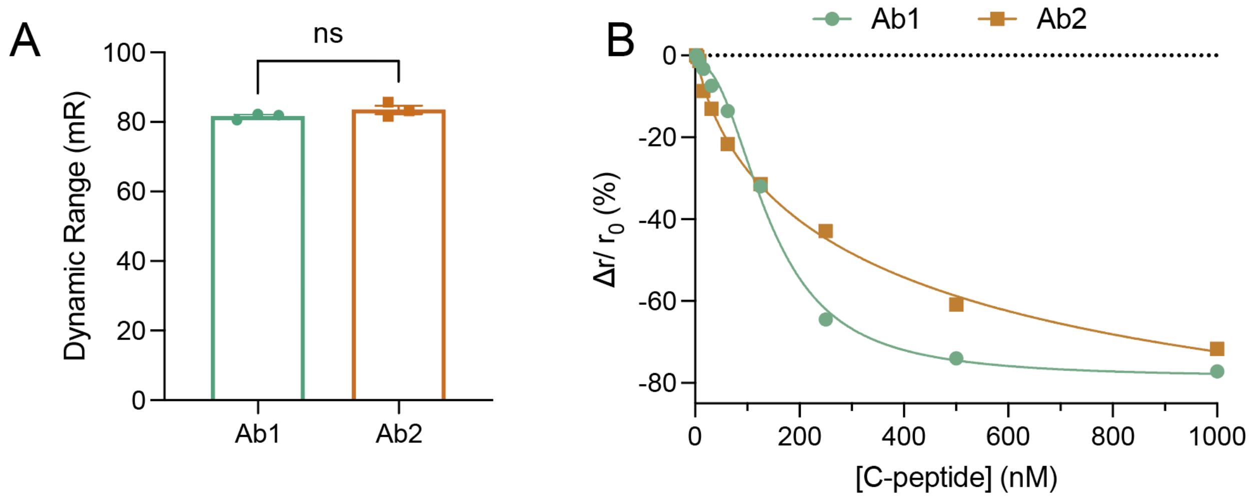


**Figure S6. Comparison of competitive binding for Ab1 and Ab2 with M6.** A) Comparison of the dynamic range of the competition curves between Ab1 + M6 and Ab2 + M6 after 5min incubation. B) Comparison of the competitive binding curves between Ab1 + M6 (**Figure 5C**) and Ab2 + M6 (**Figure 6F**) after 5-min incubation.

**Table S1.** Summary of C-peptide* developed for Ab1.

One tracer containing full-length human C-peptide (F-C-peptide*) and 6 tracers containing mutant human C-peptide fragments (M1-6) were developed for Ab1. Binding characteristics are presented for each tracer. M6 (highlighted) was selected for on-chip assay due to its high dynamic range and short time-to-reach equilibrium.

| | F-C-peptide* | M1 | M2 | M3 | M4 | M5 | M6 |
|---|---|---|---|---|---|---|---|
| Residues | 1-31 | 8-31 | 8-31 | 8-31 | 8-31 | 8-31 | 12-24 |
| Point Mutations | None | V10A, L30V | L12V, L30V | V10A, S29A, L30V | V10A, S20D L30V | V10A, S20T L30V | S20T |
| $K_{D1}$ (nM) | 2.7 ± 1.1 | 3.6 ± 0.4 | 4.0 ± 0.93 | 2.1 ± 0.43 | 1060 ± 93 | 26 ± 3.9 | 42 ± 3.6 |
| $K_{D2}$ (nM) | 3.4 ± 0.21 | 2.1 ± 1.13 | 3.4 ± 1.34 | 4.1 ± 2.2 | / | 28 ± 2.6 | 32 ± 0.36 |
| Dynamic range at equilibrium (mR) | 43 ± 0.11 | 90 ± 2.7 | 84 ± 0.69 | 93 ± 3.0 | / | 32 ± 0.13 | 82 ± 0.93 |
| Time-to-reach-equilibrium (min) | 30 - 60 | 30 - 60 | 30 - 60 | 30 - 60 | / | 1.1 | 1.2 |

**Table S2.** Summary of C-peptide* developed for Ab2.

One tracer containing full-length human C-peptide (F-C-peptide*) and 2 tracers containing mutant human C-peptide fragments (M6-7) were developed for Ab1. Binding characteristics are presented for each tracer. M6 (highlighted) was selected for on-chip assay due to its high dynamic range and short time-to-reach equilibrium.

| | F-C-peptide* | M6 | M7 |
|---|---|---|---|
| Residues | 1-31 | 12-24 | 12-24 |
| Point Mutations | None | S20T | None |
| $K_{D1}$ (nM) | 21 ± 1.1 | 70 ± 3.7 | 58 ± 1.1 |
| $K_{D2}$ (nM) | 1.6 ± 0.67 | 77 ± 3.7 | 110 ± 10 |
| Dynamic at equilibrium range (mR) | 20 ± 1.6 | 84 ± 2.1 | 58 ± 2.8 |
| Time-to-reach-equilibrium (min) | < 5 | 0.3 | 0.25 |
